# Gut Microbiome Profiling of a Rural and Urban South African Cohort Reveals Biomarkers of a Population in Lifestyle Transition

**DOI:** 10.1101/2020.02.27.964023

**Authors:** OH. Oduaran, FB. Tamburini, V. Sahibdeen, R. Brewster, FX. Gómez-Olivé, K. Kahn, SA. Norris, SM. Tollman, R. Twine, AN. Wade, RG. Wagner, Z. Lombard, AS. Bhatt, S. Hazelhurst

## Abstract

**Background:** Comparisons of traditional hunter-gatherers and pre-agricultural communities in Africa with urban and suburban Western North American and European cohorts have clearly shown that diet, lifestyle and environment are associated with gut microbiome composition. Yet, little is known about the gut microbiome composition of most African adults. South Africa comprises a richly diverse ethnolinguistic population that is experiencing an ongoing epidemiological transition and concurrent spike in the prevalence of obesity, largely attributed to a shift towards more Westernized diets and increasingly inactive lifestyle practices. To better characterize the microbiome of African adults living in more mainstream lifestyle settings and to investigate associations between the microbiome and obesity, we conducted a pilot study in two South African cohorts that are representative of urban and rural populations. The study was designed collaboratively with community leaders. As the rate of overweight and obesity is particularly high in women, we collected single time-point stool samples from 170 HIV-negative women (51 at Soweto; 119 at Bushbuckridge), performed 16S rRNA gene sequencing on these samples and compared the data to concurrently collected anthropometric data.

**Results:** We found the overall gut microbiome of our cohorts to be reflective of their ongoing epidemiological transition. Specifically, our results show a relatively higher than expected abundance of Western gut-associated taxa such as *Barnesiella* and the presence of *Bifidobacteria* and *Bacteroides* together with the more traditionally non-Western gut-associated *Prevotella*, *Treponema* and *Succinivibrio*. Interestingly, we observed a relatively higher abundance of the *Melainabacteria*, *Vampirovibrio*, a predatory bacterium, in the rural cohort. We also found *Prevotella*, despite its generally high prevalence relative to all taxa present in the cohort, to be associated with obesity.

**Conclusions:** Altogether, this work identifies putative microbial features associated with host health in a historically understudied community. Furthermore, we note the crucial role of community engagement to the success of a study in an African setting, the importance of more population-specific studies to inform targeted interventions as well as present a basic foundation for future research in this regard.

## Background

There have been relatively few studies of the human gut microbiome in Africa, with most reported studies to date focusing on the extremes of non-Western traditional hunter-gatherer and agriculturalists African populations, as well as children with nutritional deficiencies^1–4^. A consistent finding of these studies is the inverse relationship between the *Bacteroides* and *Prevotella* genera of the *Bacteroidetes* phylum. *Prevotella* is associated with plant-based diets predominantly in non-Western populations, whereas increased relative abundance of *Bacteroides* is thought to result from animal fat- and protein-based diets^5–10^. It is important to note that across most of sub-Saharan Africa, the dominant lifestyle has been agricultural for at least 1000 years^11^, with relatively few people living hunter-gatherer or pastoralist lifestyles. However, over the last 50 years in particular, the epidemiologial transition discussed throughout this paper has had significant impact on many lifestyle practices.

Other studies have focused on the role of the microbiome in areas of public health concern on the African continent, including nutrition, vaccine response efficacy, the impact of antibiotics, mental health and human immunodeficiency virus (HIV)^12–15^. Obesity, a growing health burden^16^ on the African continent, has received comparably less attention from microbiome researchers. The dramatic increase in the prevalence of obesity has been attributed, in part, to the ongoing epidemiological transition on the continent towards more Westernized practices, such as the consumption of more animal-based and processed products with increasing physical inactivity^17–19^, further complicating the existing challenge of malnutrition facing the continent^20,21^. This is reflected in an analysis of demographic and health survey data from 24 African countries^16^ where the prevalence of overweight and obesity among women increased in all 24 countries with either a doubling or tripling in the incidence of obesity reported in 50 percent of the surveyed countries. Pertinent to this study is the statistics indicating black South African women to have the highest prevalence of obesity (42%) within sub-Saharan Africa^22^ with general continental body mass index (BMI) trends showing a decline in the underweight population with a concomitant increase in the overweight and obese population^23–25^. The implication of this is the potential increase in the prevalence of comorbidities including diabetes and other cardiometabolic diseases augmenting the health and economic burden in African societies^26–28^.

Globally, several studies have focused on understanding the apparent dysbiosis observed in obesity^29,30^. Early findings suggest notable differences in the *Firmicutes*:*Bacteroidetes* ratio in lean versus obese individuals. Furthermore, lean individuals demonstrate relatively higher microbial diversity and an enrichment in microbial species associated with anti-inflammatory properties^31,32^. African populations have, however, been understudied in these efforts. Consequently, there is a paucity of data within Africa comparing the gut microbiota of obese individuals to their leaner counterparts. This is crucial, as differences in dietary and environmental exposures may render findings in non-African populations poorly generalizable to the African context, especially with the ongoing epidemiological transition in Africa^4,33,34^.

Here, we present a study that investigated the gut microbial composition of two South African cohorts with a focus on obesity. South Africa, with its diverse ethnolinguistic groups, presents a unique opportunity to study the effects of this transition on the gut microbiome. With obesity being an established risk factor in cardiometabolic diseases, understanding the within-population differences observed between obese and lean individuals in this setting could prove critical to improving our understanding of its association to the pathogenesis of the disease.

This pilot study was nested in the AWI-Gen project^35^, a part of the Human Heredity and Health in Africa (H3Africa)^36^ initiative. AWI-Gen is a collaborative effort, with participants in six sites across four African countries, established to assess genomic and environmental factors that influence cardiometabolic diseases risk, with the aim of informing treatment and intervention strategies. The study focused on characterizing the gut microbiome of lean and obese female adults from two cohorts comprising communities across two South African provinces, Gauteng and Mpumalanga, representative of relatively urban and rural lifestyles respectively. These cohorts are managed by established health and demographic surveillance sites (HDSS) in partnerships with the University of the Witwatersrand (Wits) and the Medical Research Council (MRC) of South Africa. The Agincourt HDSS^37^ in Mpumalanga encompasses a collection of rural communities in the Bushbuckridge municipality that are undergoing rapid epidemiological changes which may allow for some of the areas to be classified as peri-urban. The Developmental Pathways for Health Research Unit (DPHRU) in Gauteng, on the other hand, is focused on Soweto, a highly urbanized area that borders Johannesburg. It comprises individuals that have been urbanized for many generations even though in-migration remains at a high level.

In this study, we performed 16S rRNA gene analysis of the gut microbiome of 170 female individuals in Bushbuckridge and Soweto. Using BMI values, we assessed compositional differences in the microbiome between lean and obese individuals within and between Bushbuckridge and Soweto. The overall microbial composition of the sampled data was evaluated to improve our knowledge of the general microbiota landscape of these representative cohorts. We also provide insight into the feasibility of such studies in rural communities whilst highlighting the importance of community engagement to this effort.

## Results

### Community Engagement and Sample Collection

The research team visited the Agincourt HDSS study area (rural Bushbuckridge site) during the planning phase of the study to discuss the proposed study with the Community Advisory Group (CAG), who are now permanent members of a study Research Advisory Group (RAG). Discussions with the RAG involved potential concerns of community members and reactions to collecting stool samples, as well as how to practically collect samples. It was deemed appropriate to attempt to collect stool in this population, provided that a detailed explanation of its intended usage would be provided by the trained field workers. Specifically, the RAG communicated that it was important to convey to participants that the stool would only be used solely for the proposed study, not be sold, and that any remaining sample would be disposed of after the of the study. In addition to interactive meetings with the RAG, graphical flyers with an overview of the research aims were made available to participants. An instructional video recording was provided to assist the training of field workers. It was also agreed upon that samples will be collected from participants in a culturally sensitive manner as available research stool collection kits are designed for use on Western toilets with plastic toilet seats, and most households in the study area use pit latrines with concrete or no toilet seats.

Sixty-five (65) responses were received from a follow-up survey done on the first 100 participants in the Bushbuckridge area to get their feedback on the process, with one refusal to participate. All 65 survey participants responded positively indicating an interest in future participation. Most found the instructions clear and useful, and they had not found participating embarrassing or unpleasant. This survey was conducted telephonically, and we had a concern that the participants’ responses may not have been entirely forthright. Several participants also asked about getting results back.

Following preliminary data analysis, the research team organized an interactive workshop with 18 CAG representatives on-site. The session reiterated the importance of the study, conveyed initial results, and solicited feedback from community members. Participants were encouraged to share their insights and learnings with a “concept map,” a diagram that depicts relationships between different ideas, such as “bacteria”, “health” and “disease. This provided the research team with a sense of the communal understanding of the study.

### Participant Recruitment and Study Cohort

With ethics approval from the Human Research Ethics Committee (Medical) of the University of the Witwatersrand (M160121) and the Provincial Health Research Committee of the Province of Mpumalanga (MP2017TP22851), 132 female individuals from Bushbuckridge (28.4°E, 31.2°S) and 58 from Soweto (26.2485° S, 27.8540° E) were recruited for the study. However, only 170 participant samples (Bushbuckridge: 119, Soweto: 51) were included in the study due to confounding factors to the focus of this pilot (18 HIV-positive samples and two samples with collection irregularities were excluded). The age and BMI distribution of the cohorts are shown in Table 1.

**Table 1.** Age and BMI distribution of cohorts.

| | Mean ± SD | Median | Range |
| --- | --- | --- | --- |
| **Age (years)** |  |  |  |
| Bushbuckridge | 53.26 ± 7.88 | 53 | 41 - 70 |
| Soweto | 49.86 ± 5.50 | 50 | 40 - 59 |
| **BMI (Kg/m<sup>2</sup>)** |  |  |  |
| Bushbuckridge | 32.6 ± 7.95 | 31.17 | 21.23 - 58.97 |
| Soweto | 36.05 ± 9.23 | 36.52 | 20.43 - 58.62 |

### Pre-processing and Quality Control

16S rRNA gene sequencing was performed with primers to the V3 and V4 regions. A total of 16,595,564 sequences were obtained from the 169 samples that met the quality control criteria. The sequence depths ranged from 12 to 160,908 reads per sample (Supplementary Table 1), with a mean of 97,620.96 ± 2371.40 and a median of 96,951.5, resulting in a total of 10,640 unique amplicon sequence variants (ASVs) with redundant taxonomies. As a result of relatively low sampling depths of 12 and 25,277 reads for S002 and S015 respectively and the likelihood that the richness of the samples was not fully observed at their sequenced depths, they were further excluded from downstream analyses (Figure 1). Also excluded from further analysis was sample S017 as it failed to meet the set quality control criteria. This resulted in a minimum sequence depth of 53,587 reads for 167 samples. As expected, the taxonomies associated with the corresponding ASVs accounted for two kingdoms (*Archaea* and *Bacteria*) resulting in 17 phyla, 31 classes, 48 orders, 93 families, 255 genera and 241 species, with unclassified ASVs also detected at all levels (Table 2). These numbers represent non-redundant taxa. The number of classified species (in parenthesis), were representative of the following genera: *Bacteroides* (21), Lactobacillus (13), *Prevotella* (9), *Alistipes* (8), *Clostridium _sensu_stricto* (7), *Treponema* (3) and *Bifidobacterium* (4).

**Table 2.** Distribution of taxonomic classification of ASVs in sampled South African pilot dataset.

| Taxa Level | Classified ASVs | Associated Taxa | % Unclassified ASVs |
| --- | --- | --- | --- |
| Kingdom | 10,562 | 2 | 0.73% |
| Phylum | 10,333 | 17 | 2.89% |
| Class | 9,988 | 31 | 6.13% |
| Order | 9,879 | 48 | 7.15% |
| Family | 8,847 | 93 | 16.85% |
| Genus | 6,750 | 255 | 36.56% |
| Species | 346 | 241 | 96.75% |

**Figure 1.**
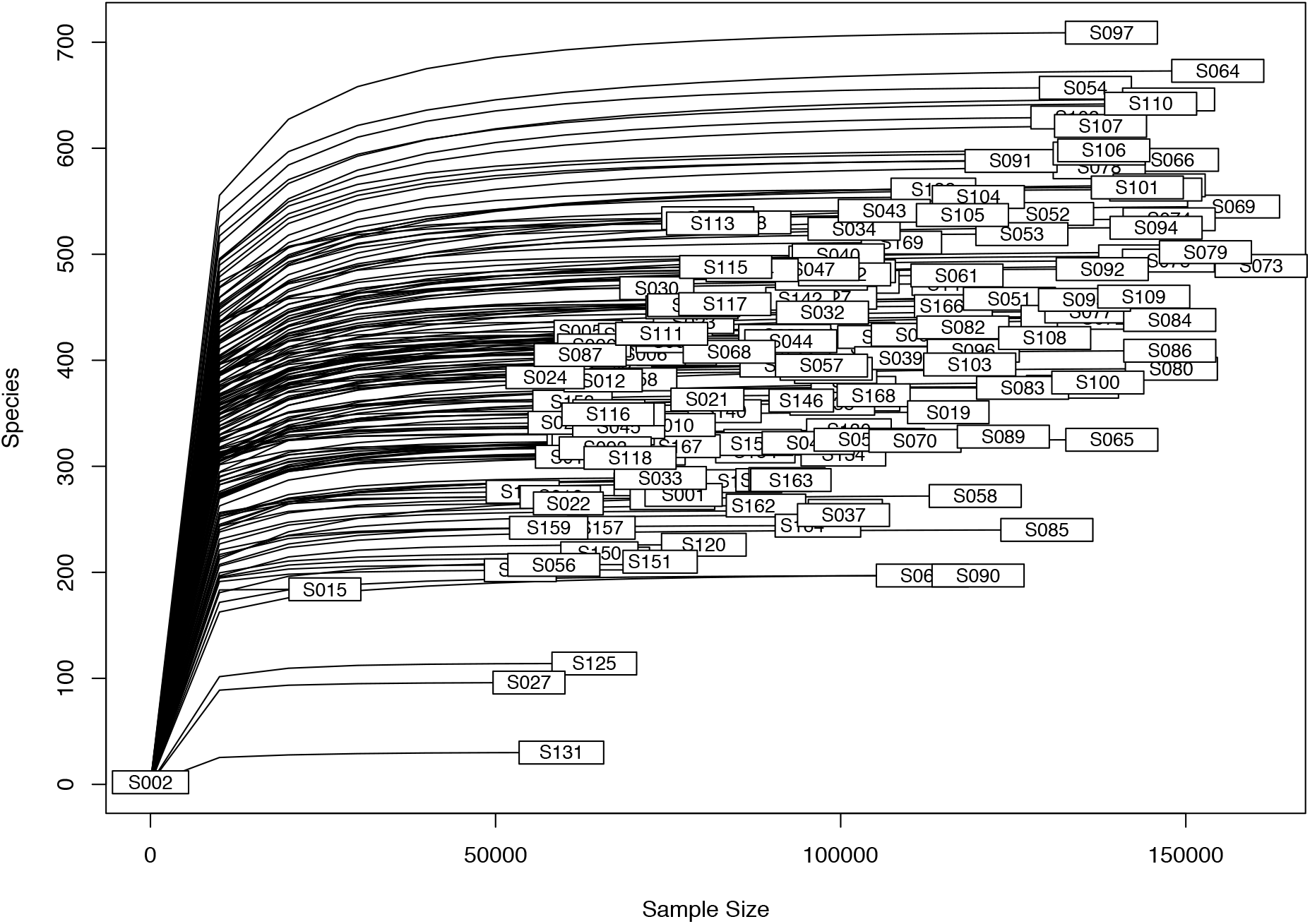
Rarefaction curve of sampled data. S002 and S015 were excluded from further analyses.

### Microbial Community Richness Estimates and Differences

With the majority of diversity metrics being sensitive to varying sequencing depths across samples^38^, rarefaction was done at a read depth of 53,500 to maximize the capture of the observed microbial taxa richness in the cohort. This cut-off was chosen based on the spread of the read depths as visualized in the rarefaction plot in Figure 1, with the exclusion of two very low-depth samples.

#### Site differences

In a cohort-wide comparison to evaluate overall differences between the Bushbuckridge and Soweto sites irrespective of BMI status, statistically significant p-values were observed for Shannon^39^ alpha diversity measure (0.01) (Figure 2) and the weighted and unweighted uniFrac^40^ beta diversity measures (< 0.001), visualized in principal coordinate analysis (PCoA)^41^ plots (Figures 3A and 3B). We find that regardless of clinical grouping (lean/obese), geographical location took precedence with respect to sample clustering. The PCoA plots also present a moving divide between rural Bushbuckridge and urban Soweto. This appears to reflect a transitional state possibly owing to gradual lifestyle and dietary changes.

**Figure 2.**
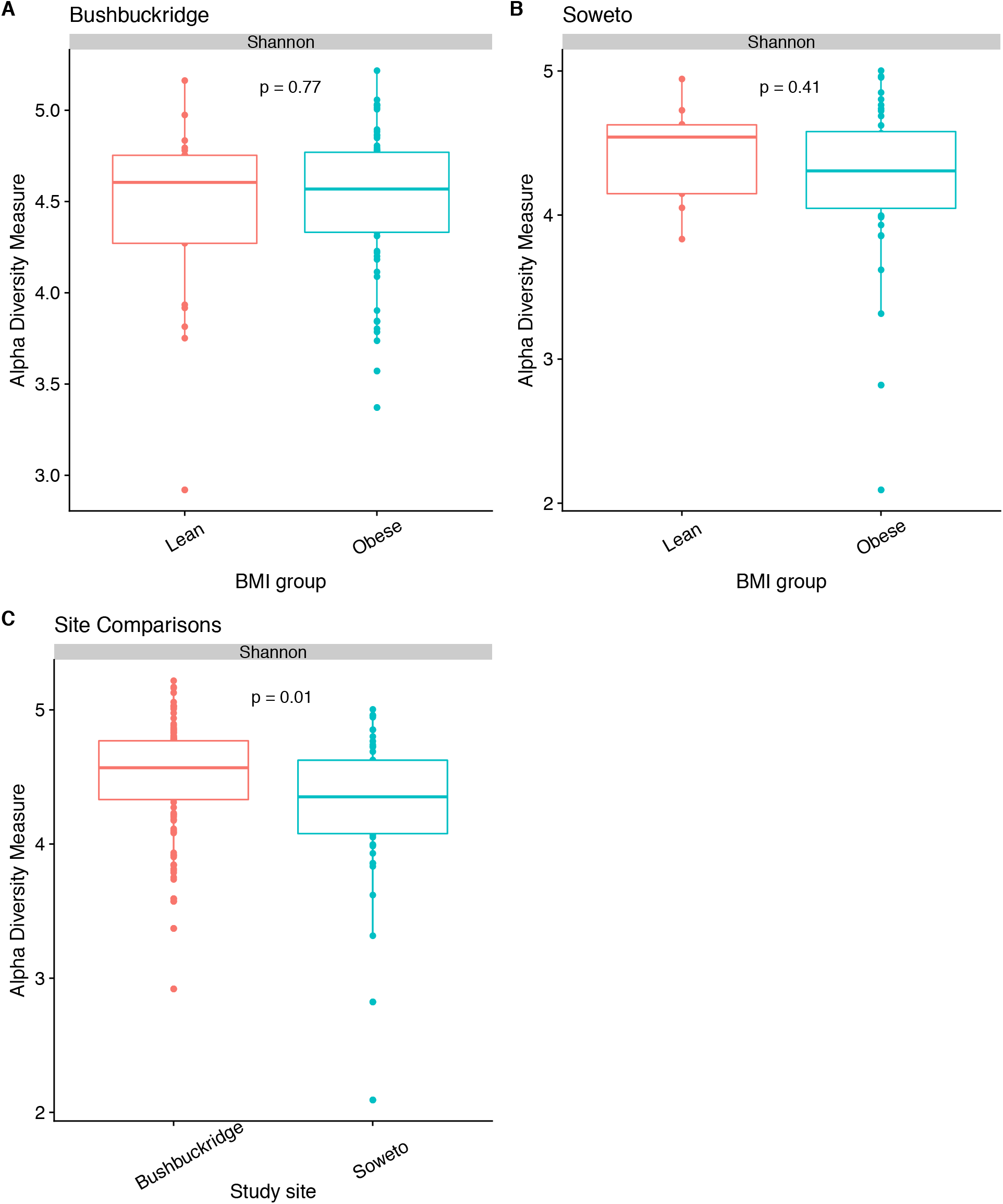
Boxplots of Shannon alpha diversity measure estimates. Site-specific alpha diversity comparisons of lean vs obese samples in (A) Bushbuckridge and (B) Soweto. Study site differences are shown in (C).

**Figure 3.**
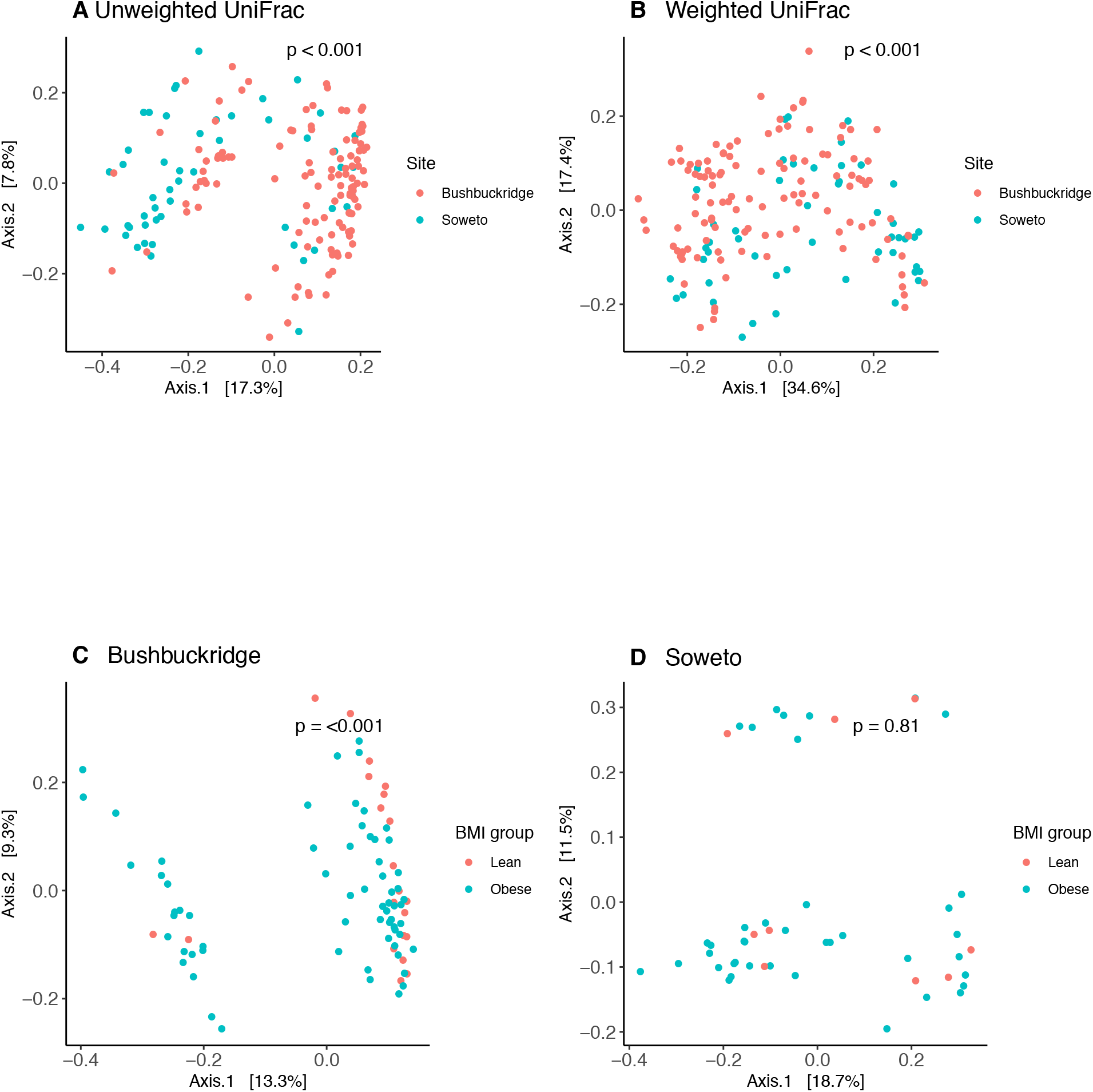
Beta diversity PCoA plots. Combined Bushbuckridge and Soweto datasets visualized using (A) Unweighted uniFrac and (B) Weighted uniFrac distance measures. Site-specific lean and obese sampled data using unweighted uniFrac distances in (C) Bushbuckridge and (D) Soweto.

#### BMI differences

In evaluating the potential diversity across BMI categories, Shannon diversity, a measure of richness and evenness, for the lean and obese groups in Bushbuckridge (Figures 2A and 2B) were 4.43 ± 0.52 and 4.51 ± 0.39, respectively. The exclusion of an apparent outlier in the Bushbuckridge lean group resulted in a Shannon index of 4.50 ± 0.39 in that group. The corresponding estimates for Soweto were 4.44 ± 0.36 (lean) and 4.25± 0.56 (obese). The differences between the lean and obese groups did not reach statistical significance as indicated by the non-parametric Wilcoxon rank sum test evaluating the Shannon diversity values between both groups (p = 0.77 and 0.41 for Bushbuckridge and Soweto respectively). Beta diversity (Figure 3), however, showed statistically significant differences between the lean and obese groups in Bushbuckridge with calculated unweighted UniFrac distances using the permutational analysis of variance (PERMANOVA) test (p < 0.001 for Bushbuckridge and p = 0.81 in Soweto (Table 3). Significant differences (p = 0.01) are also observed between the obese and lean groups in the combined cohorts dataset with unweighted uniFrac distance measurements (Supplementary Figure 1).

### Taxonomic Analyses

Overall, *Firmicutes* (43.2% ± 11.5%), *Bacteroidetes* (40.2% ± 12.1%) and *Proteobacteria* (12.4% ± 9.2%) were the dominant phyla observed in the combined gut microbiome data from these two South African cohorts (Figure 4A). In keeping with this finding, the classification of the top ten most abundant genera from all samples taken together were as *Prevotella*, *Bacteroides*, *Succinivibrio*, *Faecalibacterium*, *Ruminococcaceae* (unclassified genus), Clostridiales (unclassified genus), Oscilllibacter, Clostridium_XIVa, Roseburia and Ruminococcus (Figure 4B). A closer look at the representative urban site, Soweto (Figure 4D), showed a noticeably higher relative *Bacteroides* abundance (14.5% ± 14%) and lower *Succinivibrio* abundance (2.4% ± 6%) in comparison with Bushbuckridge at 7.8% ± 10.2% and 7.9% ± 10%, respectively (Figure 4F). Literature has repeatedly associated relatively higher *Succinivibrio* abundance with non-Western populations and a higher *Bacteroides* abundance with Western populations^9,34,42^. This has been hypothesized to be driven by diet^3^. *Alistipes*, a genus generally associated with animal-based diets^3,43,44^ comprised the top ten highly abundant genera in Soweto.

**Table 3.** Alpha and beta diversity significance of compared groups. Alpha diversity P-values were calculated with pairwise Wilcoxon rank sum test. Beta diversity UniFrac p-values were calculated with PERMANOVA.

| Sample Distribution | Group Comparisons | Alpha Diversity P-values | Weighted UniFrac P-values |
| --- | --- | --- | --- |
| Bushbuckridge and Soweto | Lean vs Obese | 0.77 | 0.41 |
| Bushbuckridge and Soweto | Lean vs Lean | 0.79 | 0.09 |
| Bushbuckridge and Soweto | Obese vs Obese | 0.008 | 0.003 |
| All samples | Bushbuckridge vs Soweto | 0.011 | < 0.001 |
| Bushbuckridge | Lean vs Obese | 0.77 | 0.74 |
| Soweto | Lean vs Obese | 0.41 | 0.45 |

**Figure 4.**
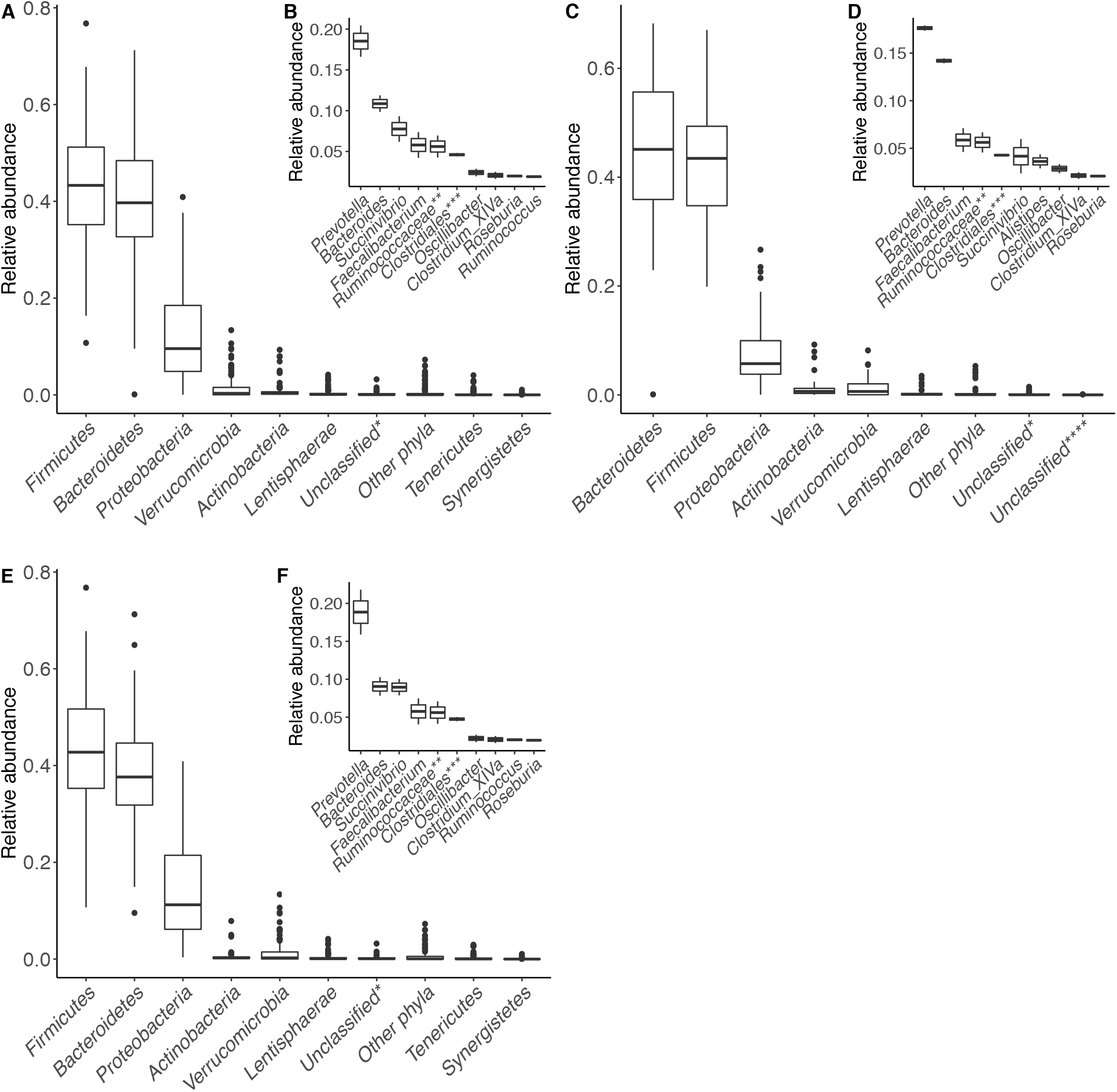
Taxonomic profiles of the gut microbiome of the sampled South African dataset. Phylum level relative abundance values are depicted in the boxplots in (A) Combined Bushbuckridge and Soweto cohorts, (C) Soweto and (E) Bushbuckridge. The inset figures B, D and F are their corresponding genera level abundance plots. “Other phyla”: phyla with mean abundance = 0. *: Unclassified bacteria phyla. **: Unclassified taxa beyond the Family level. ***: Unclassified taxa beyond the Order level. ****: Unclassified at Kingdom level.

### Microbial Compositional Analyses

To better understand the contribution of lifestyle to microbiome composition in this pilot study, the DESeq2^45^ method was applied to evaluate potential compositional differences that may not have been reflected in the alpha and beta diversity analyses. To accomplish this at site level, the data was first sub-setted to exclude the intermediate, overweight samples, while keeping only the lean (Bushbuckridge: 21, Soweto: 9) and obese samples (Bushbuckridge: 66, Soweto: 40). In addition, taxa not occurring more than two times in at least five percent of the samples were filtered out to protect against ASVs with small means and trivially large coefficients of variation across samples^46^. This resulted in a reduction in the overall observed ASVs in the cohort data from 10,640 to 1,395 ASVs representative of abundant taxa.

#### Cohort-wide Analysis

Taxonomic analysis revealed a general high prevalence of *Prevotella*. Also present in the cohorts were *Phascolarctobacterium* and *Vampirovibrio*, which were significantly higher differentially abundant in Bushbuckridge in all between-site comparisons (Figure 5). *Alistipes*, a top ten genus in the Soweto cohort in terms of relative abundance (Figure 4D), showed significantly higher differential abundance in Bushbuckridge (Figure 5C), indicating that the greater diversity in Bushbuckridge may have reduced its significance relative to other taxa in Bushbuckridge (Figure 4F). Some of the other taxa associated with Bushbuckridge include the polysaccharide-degrading *Mitsuokella* and *Dialister*^47,48^, as well as *Romboutsia*, a taxon often associated with a healthy status^49,50^, butyrate-producing *Anaerostipes*, low-fat diet-associated *Asteroleplasma*^33^ and *Alloprevotella* (Figures 5A and 5C). Soweto samples, on the other hand, showed a significant enrichment in *Bifidobacterium*, the oxalate-metabolizing *Oxalobacter*^51,52^, *Paraprevotella*, *Turicibacter*, *Peptococcus* and *Streptococcus* (Figures 5A, 5C and 6B).

**Figure 5.**
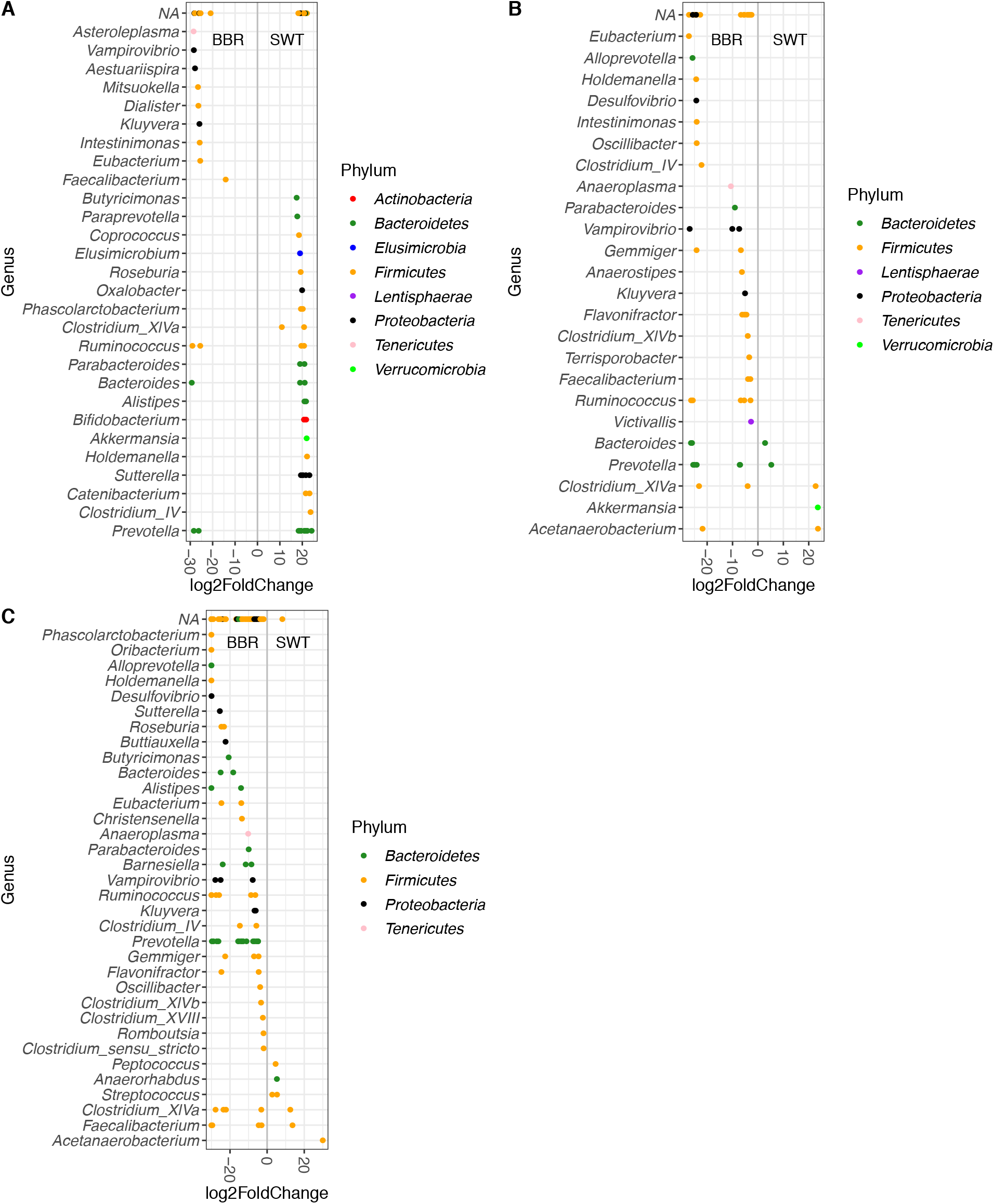
Differential abundance comparison plots of ASVs significantly abundant in Soweto (SWT) vs Bushbuckridge (BBR) in (A) Lean samples, (B) Obese samples and (C) Combined dataset – lean, overweight and obese samples. The colored dots represent ASVs differentially abundant in Soweto with respect to Bushbuckridge. Positive log2 fold change values indicate significantly enriched ASVs in Soweto and negative log2 fold change values indicate significantly enriched ASVs in Bushbuckridge. The y-axis represents the ASVs at genera level with their corresponding phyla assignments listed to the right of the plot. NA indicates ASVs that were unclassified at genera level.

Comparing the microbiomes of the combined obese group (Bushbuckridge and Soweto) with their leaner counterparts revealed *Treponema*, a previously identified hallmark taxon associated with the traditional African microbiome^7,53,54^, and *Clostridium_IV*, a butyrate producer, to be more abundant in the lean category (Figure 6C). Conversely, *Barnesiella*, *Coraliomargarita*, *Gemmiger*, *Weissella*, *Veillonella*, *Parabacteroides*, *Oscillibacter*, *Acetanobacterium* and *Alloprevotella* were among the differentially abundant taxa associated with the obese group (Figure 6C). *Prevotella*, *Bacteroides* and *Intestinimonas*, although significantly abundant across both the lean and obese groups, presented with more representative ASVs in the obese group.

**Figure 6.**
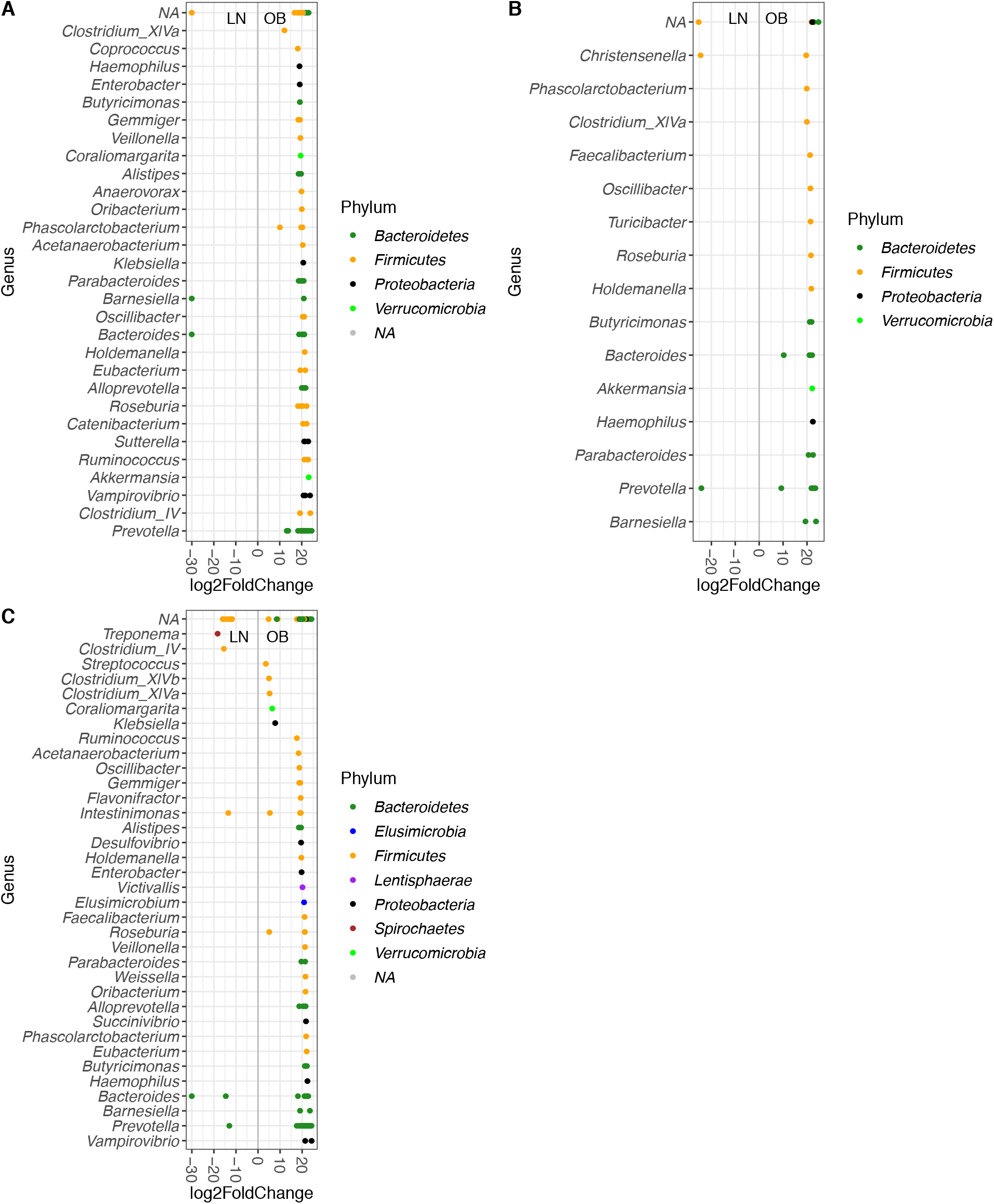
Differential abundance comparison plots of ASVs significantly abundant in obese (OB) vs lean (LN) samples in (A) Bushbuckridge, (B) Soweto and (C) Combined dataset – Busbuckridge and Soweto. The colored dots represent ASVs differentially abundant in the obese samples relative to their leaner counterparts. Positive log2 fold change values indicate significantly enriched ASVs in the obese category and negative log2 fold values indicate significantly enriched ASVs in the lean group. The y-axis represents the ASVs at genera level with their corresponding phyla assignments listed to the right of the plot. NA indicates ASVs that were unclassified at genera level.

#### Site-specific Analysis

Notably, *Prevotella* was found to be associated with obesity. This was clearly observed in Bushbuckridge, where *Prevotella* showed a higher relative abundance in the obese group. Also observed to be in higher abundance in the Bushbuckridge obese group were *Anaerovorax*, *Catenibacterium*, *Klebsiella* and *Sutterella* (Figure 6A). *Barnesiella* and *Bacteroides* were observed to be present in both the lean and obese groups (Figure 6A). In Soweto, *Prevotella* appeared to be well-represented across both lean and obese groups with a greater number of *Prevotella* ASVs present in the obese group (Figure 6B). *Haemophilus* and *Faecalibacterium* were some of the other taxa associated with the obese group in Soweto with *Christensenella* observed to have a slightly higher log2 fold change differential abundance in the lean category in Soweto. The apparent site-specific association of *Prevotella* to the obese group in Bushbuckridge is in line with literature linking the taxon to obesity^30,55,56^, although this previously reported association has contradictory results^1,2^.

### Marker Taxa Analyses

A recent meta-analysis examined differences between the gut microbial composition of traditional, rural populations and their more industrialized counterparts from several studies with datasets encompassing 13 developed or industrialized societies and two traditional hunter‐gatherer, pre‐agricultural communities^3,4,7,53,57^. The study proposed a marker taxa list distinguishing Western and non-Western bacterial communities that was corroborated by Cuesta-Zuluaga, et al^58^ by the analysis of 16 benchmark datasets with curatedMetagenomicData^59^. The dataset utilized comprised 1,655 participants from 16 countries. To further evaluate the landscape of our study data with respect to the established population-dependent compositional expectations, we compared the reported mean abundance values^58^ of Western (*Barnesiella*, *Bifidobacterium* and *Bacteroides*) and non-Western (*Treponema* and *Prevotella*) marker taxa resulting from the aforementioned analyses to their corresponding mean abundance profiles in our dataset. This was done by testing the null hypothesis that the abundances of these marker taxa were the same in the curated dataset and our sampled cohort with Student’s t-tests. Also evaluated was the null hypothesis that the mean abundances of the marker taxa in our dataset was not significantly greater than zero. Our results reject the null hypotheses for all (p < 0.002) but one taxon, *Barnesiella* with a p-value of 0.206, which presented a positive mean abundance value. We found the abundance of *Bifidobacteria*, *Bacteroides*, *Prevotella* and *Treponema*, in our data, to be intermediate between Western and non-Western microbiota, and the abundance of *Barnesiella* comparable to that in Western microbiota. This was in line with our expectations for this neither traditional nor Western dataset. However, *Barnesiella*, a Western taxon, was an exception with a higher than expected relative abundance ranging from 0.0115 to 0.0176 (Figure 7). These results reinforce the gradually changing microbial composition of the sampled cohort relative to the curated datasets.

**Figure 7.**
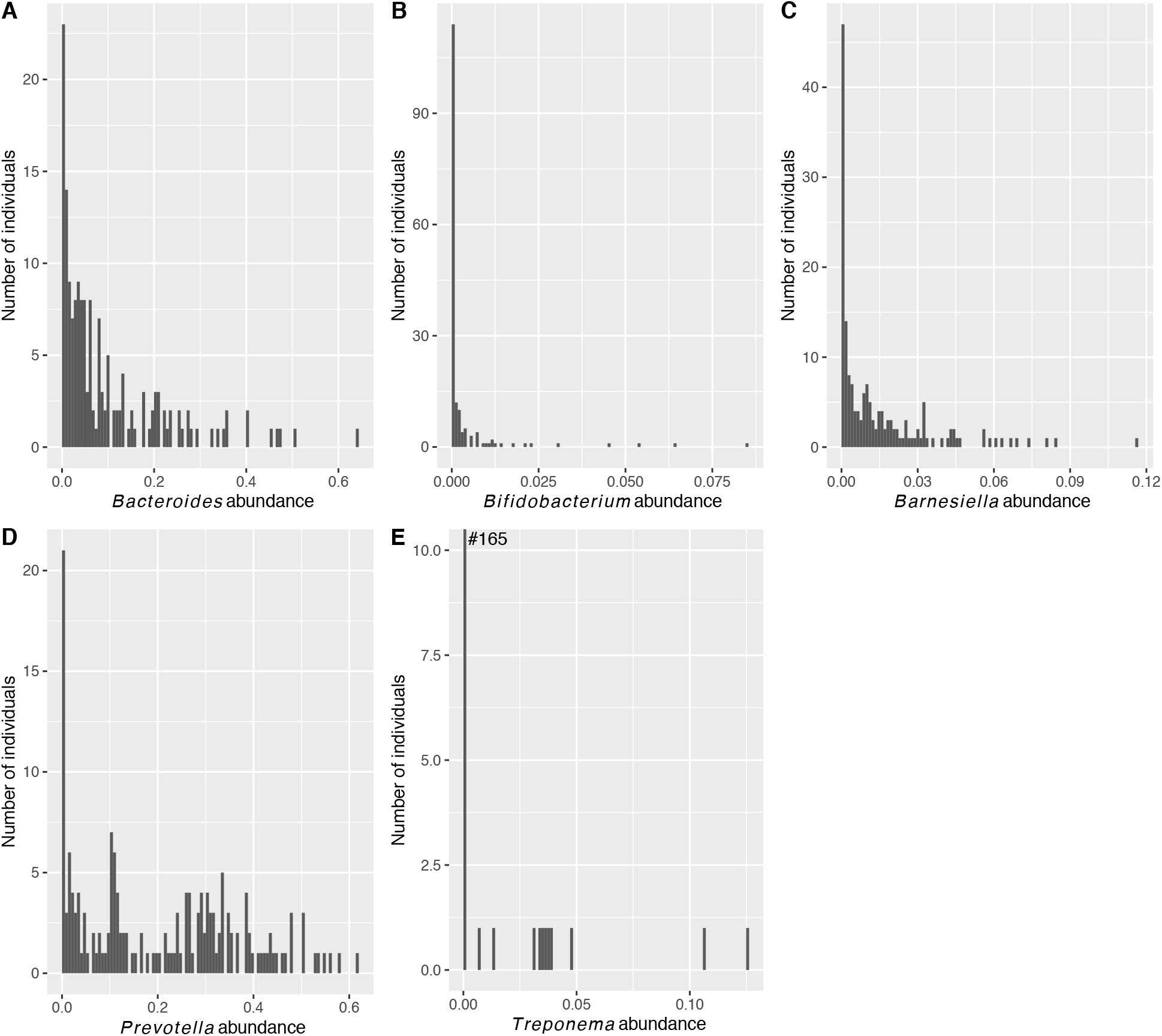
Abundances of proposed marker taxa of Western and non-Western gut microbiota in the sampled South African pilot dataset. (A) *Bacteroides* (B) *Bifidobacterium* (C) *Barnesiella* (D) *Prevotella* (E) *Treponema*. Note the scale differences between plots and the truncated axis in (E) - the number of participants with an abundance of zero is 153.

## Discussion

This study aimed to characterize the gut microbiome of two South African cohorts whilst exploring the microbial compositional differences observed in obese and lean individuals. To accomplish this, we collaboratively designed a study with active input from the community, collected stool samples and extracted DNA that subsequently underwent 16S rRNA gene sequencing. Analysis of the resulting data broadly affirms the ongoing epidemiological transitional state of our cohorts. We observed a higher than expected relative abundance of Western gut-associated marker taxa *Barnesiella* and the presence of *Bifidobacteria* and *Bacteroides*. Also present in our dataset were the more traditionally non-Western gut-associated *Prevotella*, *Treponema* and *Succinivibrio* with relative abundances comparatively lower against benchmarked marker taxa datasets. Within our cohorts, we found *Vampirovibrio*, a predatory *Melainabacteria* to present with higher relative abundances in the rural samples and *Prevotella*, despite its generally high prevalence relative to all taxa present in the cohort, to be associated with obesity. Overall, we identified putative microbial features associated with host health and highlight the importance of population-specific considerations in microbiome research. Importantly, we also shed light on the vital role of engaging the community of interest towards the success of such studies in an African setting. This study is the first on the African continent to better understand the effects of the ongoing epidemiological transition on microbial compositional changes in the context of obesity.

The study participants were recruited from two sites, about 483km (300 miles) apart that represent relatively urban and rural lifestyle and diet-practicing populations. The recruitment process and sample collection for this study relied on extensive community engagement in conjunction with a CAG at the rural site, Bushbuckridge. Although the community was familiar with the general research process, the concept of stool donation was relatively unfamiliar^60,61^. Stool collection for microbiome research purposes had never before been carried out in this population. With prevailing traditional beliefs concerning stool carrying the soul, it was crucial to be sensitive and respectful whilst clearly presenting the importance and proposed usage of the stool samples as well as the aims of the research in understandable language.

We struggled with deciding on the amount and type of information to report back, as DNA sequencing of stool is not a gold standard diagnostic test, and participants are used to receiving actionable health information as part of most studies at the HDSSs (e.g. HIV status or presence of a pathogen, like bilharzia). The HIV test, BMI and glucose level measurements done as part of the study were reported back. However, for microbiome data, science is not at the stage where meaningful results can be given, and so individual feedback would not be useful and may be potentially damaging and raise anxiety levels. This issue was raised with the RAG and understanding was shown. Nevertheless, we hope to work out a way of giving group feedback that be meaningful and if not useful at least interesting to our participants. In addition to the CAG’s valuable advisory role to the success of the collection process, some of the participants present at the feedback session noted the disposition of the fieldworker to be critical to their consenting to the study. We had a high participation rate in Bushbuckridge. Within our cohort, microbial composition reflected a transitional state comprising both Western- and non-Western-associated taxa. *Prevotella*, *Succinivibrio*, *Treponema* and the highly abundant *Ruminococcaceae* represented the traditional hunter-gatherer taxa. *Mitsuokella*, like *Treponema*, showed significant differential abundance only in the lean Bushbuckridge group. Although not a benchmarked traditional taxon, *Mitsuokella* has repeatedly been associated with non-Western diets and lifestyles^48,62^. *Phascolarctobacterium*, a propionate and acetate producer that has been shown to exert beneficial effects on its host^54,63,64^, appear to be abundant across both sites. A recent study comparing various industrialized, urban populations to traditional rural societies including Congolese Pygmies, Algerian Tuaregs, Saudi Bedouins and Amazonians from French Guiana identified *Phascolarctobacterium* to be the most significant contributing taxa to the non-Western population cluster^54^. A robust meta-analysis study that compared the gut microbiomes of urbanized and pre-agricultural populations also noted it to have relatively low abundance, and in some cases absence, in Western populations^65^.

With global research findings on the apparent dysbiosis of the gut microbiome in obesity being rather inconclusive^30–33^, we sought to evaluate the differences between obese and lean individuals within and between the two study populations. The within site differences based on alpha (Shannon) diversity were moderate and did not reach statistical significance. Similar results were obtained for beta diversity calculations with weighted uniFrac distances. However, using unweighted uniFrac distances, statistical significance was reached in all comparisons (p < 0.02) with the exception of Soweto (p = 0.81). The sensitivity of these indices may have been affected by the disproportionate sample sizes. In the differential abundance analyses, large logarithmic fold changes were observed in the component microbial taxa of the obese samples relative to their leaner counterparts. *Sutterella* and *Klebsiella*, which have been previously associated with obesity^32,66^, as well as *Alloprevotella* are among the differentially abundant taxa in the obese samples in Bushbuckridge. *Oscillibacter* was associated with cohort-wide obesity irrespective of site. The association of *Oscillibacter* to obesity has been previously reported in a European cohort^67^. Overall, the lean comparisons showed slightly greater diversity than the obese groups with taxa representative of eight different phyla (Figure 5A). The PCoA plot for Bushbuckridge (Figure 3C) demonstrates a divide between samples that do not appear to be entirely driven by BMI categories. It is, however, possible that associations with small effect sizes exist in our sampled cohort that could be detected with larger sampling. Also, as limited demographic and dietary data were collected for this pilot, further exploration is warranted.

Despite not reaching statistical significance when compared to benchmarked datasets, the large log2 fold changes observed in the abundance of *Barnesiella* in Bushbuckridge is noteworthy. The emergence to significant abundance of such an otherwise “Western” taxon highlights the evolution being observed in the microbial composition across transitional African populations. *Barnesiella* has been shown to facilitate anti-cancer immunomodulatory responses^68^ as well as being associated with the clearance of and protection from Vancomycin-resistant Enterococcus (VRE) colonization^69,70^, one of the leading causes of nosocomial infections^71^. With the seeming range of protection *Barnesiella* confers, it may play a crucial role in maintaining the homeostasis of the functional microbiome amidst ongoing changes and thus worthwhile to be explored further. Also curious in the Bushbuckridge cohort was the predatory *Vampirovibrio*. Although not very well-studied in humans to date, *Vampirovibrio* is capable of invading and attacking other bacteria without harming human cells and has been proposed for further studies in bioremediation^75^ to reduce the use of antibiotics. *Melainabacteria*, the phylum to which *Vampirovibrio* belongs^72,73^, is generally found to be present in aquatic habitats as well as associated with the guts of herbivorous mammals and humans with predominantly plant-based diets. They are also known to synthesize vitamins B and K, which in addition to their fiber-digesting abilities posits them as beneficial bacteria to their hosts. The presence and high log2 fold change value of this bacteria in Bushbuckridge relative to urban Soweto highlights the different stages of these two cohorts in the microbial compositional changes occurring with the gradual lifestyle and dietary shifts towards more Western practices. It is important to note that *Vampirovibrio* is classified under the *Proteobacteria* phylum by the Ribosomal Database Project (RDP)^74^, the database used for taxonomic classification in this study.

Several studies have identified obesity-associated taxa primarily in non-African populations^23,48,49^ despite these reported connections being inconsistent^1,21,2,67^. The differential abundance, prevalence or presence of microbial taxa across populations may require population-specific associations for relevance as universal classifications may not necessarily be generalizable. The seemingly ubiquitous presence of *Prevotella* in the sampled cohort and its association with obesity in Bushbuckridge brings to the fore the role of some *Prevotella* strains as potential pathobionts involved in various human diseases by the promotion of chronic inflammation^75,76^. Increased abundance of *Prevotella* species at mucosal sites have been linked to several diseases including metabolic disorders and low-grade systemic inflammation^30,55,77^, a feature associated with obesity. *Prevotella*, may thus present as a critical taxon in the obesity pandemic on the African continent. Further in-depth studies to ascertain the influence of its prevalence in a community undergoing such epidemiological transition will be insightful as the beneficial or detrimental effects of *Prevotella* may very likely be dependent on its interaction with the prevailing lifestyle and environment.

## Conclusions

It is important to note that although this study may have been limited by uneven sampling, it is a pilot study that provides us with a foundation to inform future studies and highlights the vital role of community engagement in planning such studies in an African setting. A clear outcome of this study was the between-site differences across all categorical and statistical comparisons (with the exception of the lean group), with the rural Bushbuckridge site harboring relatively more diverse microbiota than the urban Soweto site. In broad summary, the compositional taxa of the gut microbiome of the representative urban and rural ethnolinguistic groups are reflective of an epidemiological transitional state and the beneficial or detrimental effects of *Prevotella* are very likely diet- and lifestyle-dependent. Lastly, the generally lower mean abundances of the proposed Western and non-Western distinguishing marker taxa in our data set in comparison with benchmarked datasets substantiates the transitional state of our African cohorts with potential implications for disease pathogenesis and general health status. Further studies with a larger sampled cohort will be very informative in this regard.

## Materials and Methods

### Community Engagement

The research team engaged the community in two interactive sessions during this study - the planning phase and post-preliminary analyses on the data resulting from the collected stool samples. A survey was also conducted on the first 100 participants in Bushbuckridge to get their feedback on the process. Prior to the collection of stool samples for the study, there was interaction with the community in conjunction with a CAG at the Agincourt HDSS (Bushbuckridge), the rural site, which gave input into the process to ensure that sample collection methods were sensitive to the community beliefs and applicable to the existing toilet facilities in the area. This group comprised eight community representatives and indunas. The meeting discussions were focused creating awareness on what the project entailed and the importance of such research in the community, as well as on potential concerns and reactions of community members to stool sample collection and the practicality of such endeavor. Also deliberated on was the role of the trained fieldworker in the recruitment process and the available resources (graphical flyers) to clearly communicate the study aims and usage of the collected stool samples in understandable language to potential participants.

The interactive workshop that followed the preliminary data analysis aimed to reiterate the importance of the study, broadly present some of the initial results and very importantly, solicit feedback from the community members and participants. As this was a pilot study, it was important to the research team to gauge the level of understanding of the study post-completion in order to inform future studies in this regard.

### Recruitment and Study Cohort

This study is nested in the AWI-Gen project, which is a part of the Human, Heredity and Health in Africa (H3Africa) consortium. AWI-Gen explores genetic and environmental factors in cardiometabolic disorders in African populations with six sites across four countries. The recruitment of participants for this study was done at two of the South African sites – the Bushbuckridge area within the Agincourt HDSS, Mpumalanga (rural) and Soweto, Johannesburg, Gauteng (urban).

Participants were randomly selected from the AWI-Gen cohort within the BMI strata defined below and are in the age range of 40 – 70years (Table 1). To minimize confounding effects, male and HIV+ participants were excluded. Participants were divided into three groups based on their BMI values – lean, overweight and obese. The lean group comprised participants with BMI < 25, the overweight group comprised participants with 25 ≤ BMI < 30 and the obese group had BMI ≥ 30. Anthropometric (height and weight) and blood pressure measurements were taken at the time of collection, and a rapid HIV test done. We also had extensive other data about participants from previous engagements. The study was approved by the Human Research Ethics Committee (Medical) of the University of the Witwatersrand (M160121) and the Provincial Health Research Committee of the Province of Mpumalanga (MP2017TP22851).

To facilitate the participant recruitment and sample collection processes, comprehensive information sessions were held with the fieldworker on the study aims and its importance. This was crucial as the recruitment success could be reliant on the fieldworker’s ability to effectively communicate these to prospective participants. The fieldworker was also aided by training videos and experience gained from self-collecting personal stool samples to facilitate relatability to the collection process.

### Sample Collection

Stool samples were collected from consented participants using DNA Genotek®’s OMNIgene microbial collection and stabilization kit and sent to the laboratory. The stool samples were subsequently aliquoted into cryovials and frozen at –80 degrees Celsius prior to DNA extraction.

### DNA Extraction and Sequencing

Frozen stool samples were thawed on ice. Genomic (total) DNA was extracted using Qiagen®’s QIAmp Powerfecal DNA kit and sent to a dedicated core facility for the sequencing of the V3 – V4 hypervariable region of the 16S rRNA gene on the Illumina MiSeq® platform using 341F 5’-CCTACGGGNGGCWGCAG-3’ and 805R 5’-GACTACHVGGGTATCTAATCC-3’ as forward and reverse primers respectively^78^.

### Sequence Data Analyses

The DADA2 (v1.10.1) pipeline^79^ was used for pre-processing and performing quality control on the sequences. Briefly, the demultiplexed paired-end sequences were imported into DADA2. Based on the quality plots, the sequences were filtered with a maximum of expected errors of 2 and 4, and sequence lengths of 280 and 240 bases for the forward and reverse reads, respectively, with primers trimmed accordingly. The resulting reads were dereplicated and merged to obtain the full denoised sequence which was used in the creation of a sequence table containing the abundance values of sequence variants from the sampled data. Chimeras were subsequently removed, and the non-chimeric sequence table was utilized for downstream analyses.

### Taxonomic Classification

The DADA2 implementation of the naïve Bayesian classifier method was applied in the assignment of taxonomies to the amplicon sequence variants using the *RDP trainset 16* DADA2-formatted reference set from the Ribosomal Database Project (RDP)^74^ and a minimum bootstrapping parameter of 50.

### Phylogenetic Tree Construction

The DECIPHER R package was utilized in performing a multiple alignment of the ASVs with the phangorn^80^ package constructing a tree based on the aligned data. Midpoint rooting was subsequently applied to the phylogenetic tree which was used to facilitate downstream diversity analyses.

### Alpha and Beta Diversity Analyses

The ASVs together with the resulting rooted phylogenetic tree, taxonomic data and sample metadata were imported into phyloseq^46^ for diversity analysis. Based on the output from the pre-processing step, rarefaction was applied at a sampling read depth of 53,500 to allow for adequate capture of the observed microbial taxa richness in the cohort as diversity metrics are generally sensitive to sample read depths.

First, Shannon^39^ alpha diversity estimates for the samples were calculated. This measure was applied to a pairwise Wilcoxon rank sum (Mann-Whitney) test to assess whether the observed ASVs differed significantly (p < 0.05) between specified categories. Boxplots were generated to visualize the categorical differences based on the Shannon diversity values. Comparisons were done as indicated in Table 4.

**Table 4.**
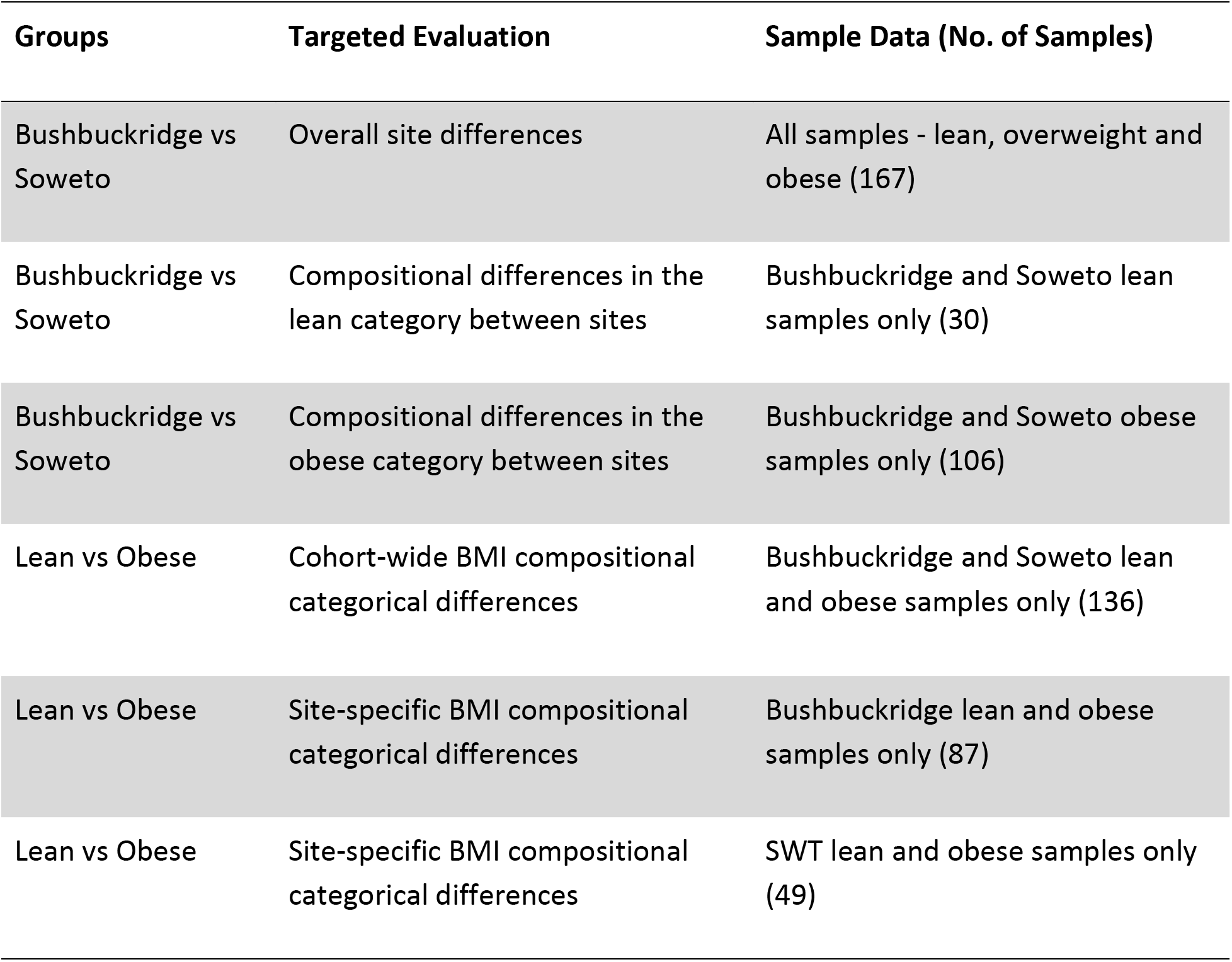
Group comparisons evaluated in this study.

Next, beta diversity between the samples was evaluated using weighted and unweighted UniFrac^40^ distance matrices for PCoA^41^ to generate relevant ordination plots. PERMANOVA analysis was done to test for differences between specified categories (Table 4).

### Differential Abundance Analyses

To evaluate differences in bacterial taxa abundance across BMI categories and sites, a negative binomial generalized linear model (DESeq2)^45^ was used. Briefly, raw counts were modelled with a negative binomial distribution and internal adjustment done for “size factors”. This adjustment normalized for differences in sequencing depth between samples. Prior to analyses, the data was filtered to exclude taxa that is not observed more than two times in more than five percent of the 167 samples. This cut-off was chosen with respect to the sample size and the general data sparsity to protect against ASVs with small mean and trivially large coefficients of variation across samples. This resulted in 1,395 high abundance ASVs being included in this analysis. DESeq2 models were adjusted for batch effects, where applicable, and BMI for the overall site analysis. Statistical significance was determined by the Wald’s test with Benjamini-Hochberg corrected p-values.

### Marker Taxa Analyses

To establish the status of our sampled cohorts along the continuum of westernization, we sought to compare the mean relative abundances of proposed Western and non-Western marker taxa as compiled by a recent meta-analysis^65^ with the corresponding values in our dataset. Mean relative abundances of these Western (*Barnesiella*: 0.0126, *Bifidobacterium*: 0.074 and *Bacteroides*: 0.230) and non-Western (*Treponema*: 0.021 and *Prevotella*: 0.245) marker taxa were obtained from the results of the analysis of 16 benchmarked datasets with curatedMetagenomicData^59^ by de la Cuesta-Zuluaga, et al^58^. Student’s t-tests were performed to test the null hypotheses that the mean abundances of these taxa are the same in both datasets and not significantly greater than zero in our sampled cohort. The proposed taxa can be used as markers of lifestyle and geographical origin in the chosen public datasets as well as in the South African cohorts.

### Feedback from Participants

The follow-up survey was done on the first 100 participants at Bushbuckridge about 3 months after collection. The survey was conducted telephonically – each person was phoned at least three times. One person refused to participate, and 65 people agreed. Participants were asked three questions on the Likert scale (1=strongly agree, 5=strongly disagree). Questions and results were: Did you find the instructions clear (1.15), did you find the instructions useful (1.46), did you find participation unpleasant or embarrassing (3.98).

## Declarations

### Ethics approval and consent to participate

The study was approved by the Human Research Ethics Committee (Medical) of the University of the Witwatersrand (M160121) and the Provincial Health Research Committee of the Province of Mpumalanga (MP2017TP22851). Consent was obtained from all study participants before any sample collection was done.

### Consent for Publication

Not applicable.

### Availability of Data and Material

The microbiome nucleotide sequence datasets generated during and/or analyzed during the current study are currently being submitted to ENA. The corresponding phenotype data is being submitted to the EGA in terms of the data sharing policy of the Human Heredity and Health in Africa consortium (H3A), the university ethics committee and consents given by participants.

Researchers may apply to the independent H3A Data and Biospecimens Access Committee to access the data which will consider each case in terms of H3A policies and to protect participants data. We expect to have persistent IDs for the data sets by the time review is complete.

## Authors’ Contributions

OHO analyzed and interpreted the data, drafted the paper and formed part of the team that extracted the DNA from stool samples with FBT and VS; FBT, VS and RB contributed substantially to the revision of the manuscript; SMT, ZL, SH, ASB conceptualized and received seed funding, and together with FXG, KK, SAN, ANW, RGW and OHO planned the project; RT facilitated the community engagement sessions at the Agincourt HDSS and contributed to the manuscript; ASB and SH contributed substantively towards the design and management of the project as well as extensively reviewed and contributed to the manuscript. All authors read through the manuscript prior to its submission.

## Acknowledgements

We thank our participants who generously gave of their time in support of African science. We relied on many people to make this happen, including Yusuf Ismail, Floidy Wafawanaka, Melody Mabuza, Michaella Hulley, Amanda Haye, and Daniel Ohene-Kwofie. We thank Michèle Ramsay for her wise counsel and leadership of the AWI-Gen consortium, DNA Genotek for the initial donation of some sample collection kits and Whitehead Scientific for timely assistance with the delivery of kits and accessory items. We are also to grateful to Dylan Maghini for the critical feedback on the manuscript.

