## Supplementary figures and images for "Gut Microbiome Profiling of a Rural and Urban South African Cohort Reveals Biomarkers of a Population in Lifestyle Transition"

### Supplementary Figure 1

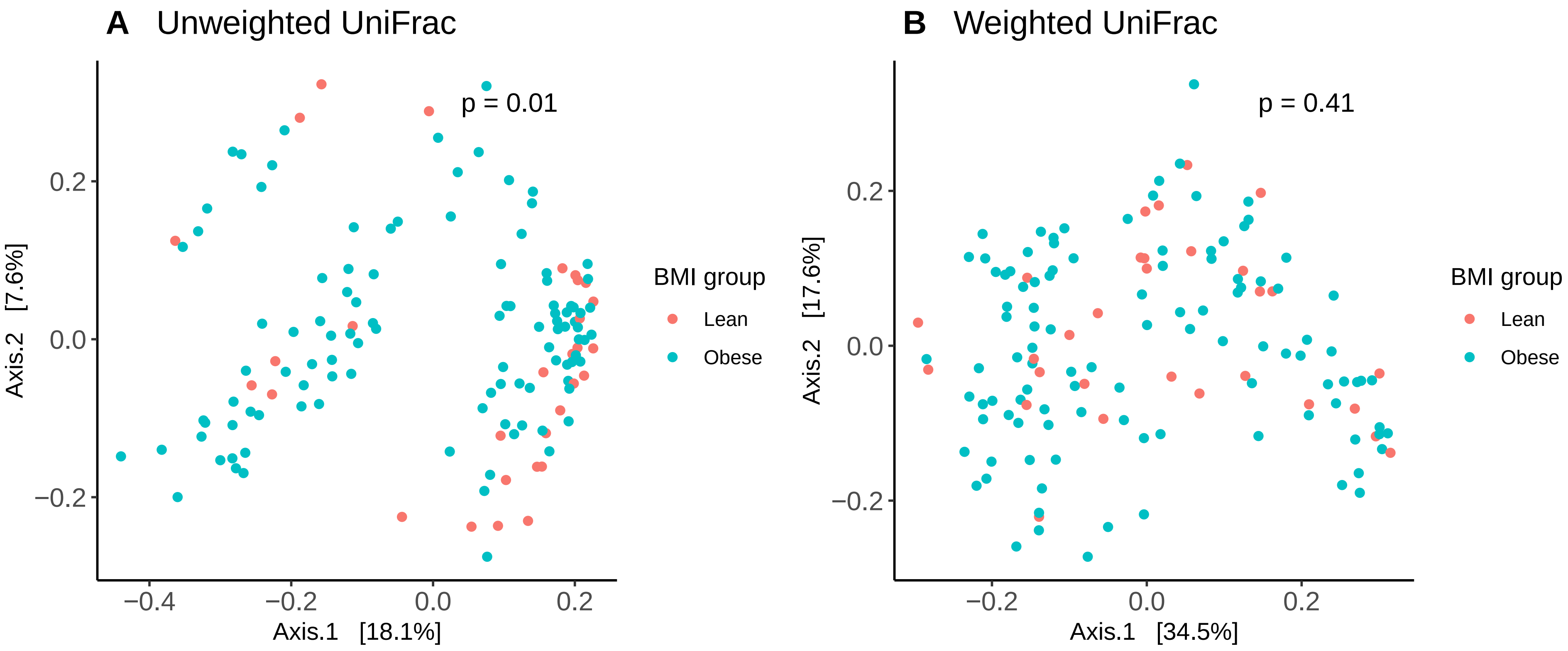
