## Supplementary Table 1 for "Gut Microbiome Profiling of a Rural and Urban South African Cohort Reveals Biomarkers of a Population in Lifestyle Transition"

| SampleID | input | filtered | denoisedF | denoisedR | merged | non-chim |
|---|---|---|---|---|---|---|
| FQOSV | 221 | 36 | 3 | 1 | 0 | 0 |
| BZEHG | 261 | 36 | 12 | 12 | 12 | 12 |
| ELHZE | 59249 | 30367 | 29106 | 29244 | 25916 | 25277 |
| DZPJR | 78941 | 64448 | 63486 | 63807 | 60682 | 53587 |
| CSK0E | 87888 | 66289 | 65163 | 65465 | 61943 | 53929 |
| JQKFV | 79201 | 63393 | 62819 | 62924 | 60297 | 54842 |
| HPRXQ | 79678 | 64065 | 61474 | 62984 | 58474 | 57178 |
| CXCOE | 95563 | 71776 | 70519 | 71025 | 67619 | 57691 |
| MB17034 | 86337 | 66140 | 65011 | 65556 | 63300 | 58441 |
| EZWQX | 90765 | 73030 | 71515 | 72248 | 67544 | 59374 |
| 3602219 | 97310 | 72774 | 72482 | 72417 | 71377 | 59537 |
| HVFJZ | 87522 | 71140 | 68757 | 69905 | 64787 | 59758 |
| HFZEJ | 84861 | 69454 | 68302 | 68755 | 65468 | 60540 |
| CRCOS | 99531 | 74434 | 72268 | 73445 | 68491 | 61134 |
| EKZCV | 84087 | 69699 | 68316 | 68855 | 65209 | 61144 |
| BKHRQ | 90134 | 73819 | 70597 | 72356 | 65697 | 61553 |
| MB17068 | 96090 | 73955 | 72057 | 73056 | 68175 | 62250 |
| CTBOZ | 90932 | 73767 | 71923 | 72802 | 68185 | 62901 |
| CBSNC | 88959 | 72609 | 69967 | 71275 | 66010 | 63301 |
| CLLYS | 89135 | 72568 | 69636 | 70990 | 65539 | 63419 |
| EDFHQ | 89675 | 72829 | 70835 | 71757 | 67634 | 63783 |
| JLNGC | 89603 | 74552 | 71821 | 73378 | 68359 | 64285 |
| 2123689 | 104000 | 80287 | 79441 | 79661 | 77775 | 64305 |
| CKXOB | 101589 | 77480 | 76155 | 76580 | 72867 | 65043 |
| CWIOI | 102654 | 76057 | 74335 | 75095 | 71233 | 65313 |
| DZWHL | 91654 | 74241 | 71333 | 73081 | 67578 | 65580 |
| MB17074 | 102521 | 78913 | 77184 | 77992 | 73978 | 65885 |
| 3379474 | 101349 | 77862 | 76854 | 77227 | 74333 | 66176 |
| MB17104 | 104096 | 80173 | 78677 | 79245 | 74443 | 66280 |
| KDRZJ | 91677 | 75529 | 72950 | 74254 | 69610 | 66412 |
| 4152135 | 102381 | 75777 | 72611 | 74391 | 68648 | 66495 |
| JRWPH | 96148 | 77928 | 75022 | 76776 | 70819 | 67357 |
| MB17019 | 111382 | 84845 | 82946 | 83851 | 78563 | 67841 |
| MB17100 | 106172 | 82120 | 80411 | 81139 | 77518 | 67868 |
| MB17106 | 106678 | 77943 | 76122 | 77109 | 73872 | 69487 |
| CWQ0Q | 109737 | 80976 | 78334 | 79787 | 74207 | 70278 |
| HDWHX | 103715 | 84307 | 81271 | 83002 | 77463 | 70843 |
| DFEZM | 100008 | 82292 | 80462 | 81373 | 76555 | 71064 |
| CQMOC | 112109 | 85945 | 84138 | 85064 | 79963 | 71419 |
| CQGFO | 100291 | 80491 | 77143 | 79143 | 73358 | 71662 |
| CCPOB | 104275 | 78025 | 76095 | 77116 | 73668 | 72783 |
| 1186710 | 119766 | 90607 | 89386 | 89805 | 85320 | 73306 |
| KLHYD | 105653 | 85671 | 80965 | 83916 | 75536 | 73360 |
| CLNOT | 116654 | 90123 | 88505 | 89186 | 85162 | 73842 |
| MB17006 | 114094 | 85285 | 83863 | 84421 | 80679 | 73864 |

| | | | | | | |
|---|---|---|---|---|---|---|
| MB17098 | 110346 | 84152 | 81699 | 82852 | 78000 | 74062 |
| DKWHW | 107960 | 87380 | 84519 | 86112 | 79581 | 75536 |
| 1499095 | 121348 | 92382 | 91088 | 91614 | 88092 | 75628 |
| DGXFV | 109228 | 88874 | 86096 | 87545 | 81666 | 75882 |
| CCLEW | 105355 | 85767 | 82947 | 84356 | 79151 | 76237 |
| DGEOC | 117073 | 90536 | 88719 | 89581 | 83975 | 76717 |
| BQNHZ | 106631 | 88504 | 86239 | 87334 | 81436 | 77242 |
| MB17015 | 118449 | 89128 | 85960 | 87582 | 81420 | 78455 |
| MB17025 | 122457 | 91513 | 88416 | 90117 | 84394 | 78698 |
| HGRSG | 115600 | 91005 | 87442 | 89506 | 82601 | 78701 |
| 1333396 | 119574 | 91481 | 90119 | 90844 | 87631 | 80172 |
| 2146745 | 131577 | 99844 | 96980 | 98393 | 90605 | 80251 |
| HEJHE | 114424 | 92662 | 90833 | 91291 | 87223 | 80689 |
| MB17037 | 124490 | 94039 | 90774 | 92604 | 85648 | 80738 |
| MB17101 | 128797 | 95180 | 92187 | 93871 | 87953 | 81435 |
| CAX0E | 131406 | 95917 | 92997 | 94400 | 87757 | 83188 |
| MB17105 | 124277 | 95266 | 92583 | 93906 | 87832 | 83215 |
| MB17103 | 128835 | 97664 | 94805 | 96260 | 88908 | 83338 |
| MB17046 | 128297 | 95200 | 92686 | 93918 | 88507 | 83846 |
| FRGPC | 118775 | 96739 | 93412 | 95559 | 88702 | 84571 |
| MB17010 | 130088 | 97397 | 94470 | 95817 | 88820 | 84868 |
| 1597674 | 132102 | 99833 | 97303 | 98413 | 92288 | 85331 |
| MB17069 | 128834 | 97202 | 94339 | 95928 | 90193 | 86110 |
| MB17102 | 137947 | 106472 | 103881 | 105156 | 98299 | 87170 |
| CVQON | 136122 | 103793 | 101277 | 102458 | 95897 | 87233 |
| CRXON | 134707 | 99724 | 97331 | 98504 | 92976 | 87634 |
| CKQOT | 142884 | 107801 | 105039 | 106493 | 99357 | 88692 |
| DCGOV | 136801 | 106880 | 104758 | 105666 | 100430 | 89176 |
| CDKOY | 142448 | 108652 | 106926 | 107565 | 103009 | 90320 |
| MB17012 | 141069 | 107215 | 103831 | 105633 | 97223 | 91896 |
| MB17026 | 140603 | 104737 | 102187 | 103561 | 97523 | 92075 |
| CHPOL | 142979 | 109557 | 107628 | 108492 | 102763 | 92222 |
| MB17018 | 142340 | 106160 | 103048 | 104569 | 97627 | 92773 |
| DDPOG | 140812 | 109200 | 107405 | 108071 | 103694 | 92780 |
| 3924929 | 146579 | 111883 | 109131 | 110615 | 104222 | 92938 |
| CBZOI | 146937 | 109075 | 105259 | 107364 | 99842 | 94124 |
| CFI0B | 140151 | 106963 | 103872 | 105522 | 99305 | 94289 |
| MB17020 | 146433 | 110270 | 108238 | 109146 | 103180 | 95284 |
| MB17022 | 144569 | 108789 | 105201 | 107216 | 100160 | 95992 |
| DDWOM | 146125 | 113430 | 111750 | 112353 | 107332 | 96700 |
| MB17009 | 144425 | 110417 | 107122 | 108781 | 102121 | 97203 |
| MB17035 | 139597 | 113419 | 111294 | 112296 | 106456 | 97217 |
| MB17004 | 145376 | 111605 | 108800 | 110268 | 103976 | 97359 |
| DBT0G | 146210 | 110350 | 106770 | 108642 | 101782 | 97875 |
| MB17024 | 142795 | 109827 | 107265 | 108417 | 102793 | 97896 |
| 2554637a | 148496 | 112600 | 109169 | 110889 | 103013 | 98313 |

| | | | | | | |
|---|---|---|---|---|---|---|
| 4971044 | 149527 | 114331 | 111270 | 112694 | 104911 | 98784 |
| CAWOD | 146133 | 112904 | 109923 | 111545 | 105327 | 99272 |
| MB17014 | 145991 | 113746 | 110981 | 112405 | 105849 | 99654 |
| 3956441 | 157781 | 121406 | 119254 | 120062 | 114288 | 100389 |
| MB17011 | 155730 | 123253 | 121522 | 122152 | 116162 | 100409 |
| MB17016 | 153942 | 114238 | 110402 | 112481 | 105770 | 100564 |
| CCJOV | 162002 | 122143 | 120191 | 120780 | 114928 | 100671 |
| 3480141 | 157012 | 123682 | 121779 | 122718 | 115725 | 101190 |
| 2031518 | 158401 | 120513 | 117337 | 119094 | 111183 | 101841 |
| MB17007 | 153383 | 116189 | 113175 | 114680 | 108514 | 101939 |
| CFVOM | 155322 | 115806 | 112363 | 114074 | 107144 | 102163 |
| 2266332 | 161177 | 120753 | 116710 | 118794 | 110455 | 102330 |
| 1930043 | 155389 | 117809 | 114856 | 116229 | 109581 | 102429 |
| DDZ0Q | 158181 | 120623 | 117591 | 119320 | 112443 | 102482 |
| MB17033 | 148036 | 120875 | 118966 | 119901 | 114478 | 102812 |
| 4233802 | 168833 | 129456 | 126635 | 127795 | 120407 | 104534 |
| DGV05 | 163368 | 125172 | 121939 | 123303 | 115445 | 104773 |
| MB17003 | 159836 | 124555 | 121814 | 123226 | 116916 | 106237 |
| MB17017 | 158388 | 119919 | 115906 | 118027 | 110651 | 106319 |
| DBGOS | 161666 | 123930 | 121262 | 122501 | 115806 | 106379 |
| MB17013 | 164880 | 121323 | 118591 | 119759 | 113219 | 108689 |
| DHHOH | 163885 | 124941 | 120880 | 123116 | 114785 | 108801 |
| 4858889 | 167263 | 125016 | 122105 | 123509 | 116710 | 109060 |
| MB17049 | 157620 | 130289 | 128646 | 129319 | 124154 | 110763 |
| MB17040 | 164417 | 134310 | 132121 | 132938 | 126520 | 111098 |
| MB17038 | 167023 | 135972 | 134490 | 134987 | 129166 | 111831 |
| 3663723 | 177608 | 130926 | 126675 | 129078 | 119995 | 113424 |
| GWBHF | 158344 | 126620 | 124232 | 125235 | 120598 | 115590 |
| CBY0H | 172399 | 132833 | 129388 | 131118 | 123361 | 115705 |
| DEROK | 176767 | 134015 | 130817 | 132545 | 124780 | 116328 |
| MB17039 | 170703 | 136590 | 133521 | 135099 | 127342 | 116848 |
| MB17091 | 162502 | 133574 | 130742 | 131995 | 124303 | 117640 |
| MB17062 | 163607 | 133148 | 130557 | 131514 | 124372 | 117704 |
| MB17085 | 171624 | 140787 | 138056 | 139075 | 131665 | 118695 |
| MB17077 | 170345 | 140269 | 137420 | 138855 | 130720 | 119210 |
| MB17036 | 174859 | 140985 | 138969 | 139590 | 134079 | 119494 |
| MB17071 | 174714 | 144051 | 142362 | 142785 | 137223 | 119903 |
| MB17088 | 173673 | 142720 | 139621 | 141275 | 132091 | 119943 |
| MB17070 | 169222 | 139906 | 138399 | 138795 | 133188 | 123563 |
| MB17027 | 177523 | 142703 | 139570 | 141141 | 133447 | 124444 |
| MB17072 | 174424 | 141824 | 138722 | 140185 | 133276 | 124682 |
| MB17030 | 177181 | 141767 | 138180 | 139797 | 131601 | 126249 |
| MB17063 | 181513 | 148514 | 146214 | 147024 | 139537 | 126355 |
| MB17094 | 180477 | 146320 | 143121 | 144508 | 136837 | 129553 |
| MB17066 | 190850 | 155942 | 154066 | 154707 | 147247 | 129869 |
| MB17028 | 185251 | 149853 | 146276 | 147888 | 139126 | 129976 |

| | | | | | | |
|---|---|---|---|---|---|---|
| MB17050 | 194435 | 158183 | 153720 | 156088 | 143839 | 132799 |
| MB17055 | 196883 | 160365 | 158258 | 159219 | 151513 | 133585 |
| MB17083 | 189949 | 153117 | 148986 | 150967 | 141509 | 134225 |
| MB17076 | 187526 | 152392 | 149365 | 150663 | 143340 | 135322 |
| MB17031 | 188405 | 152874 | 147561 | 150627 | 139674 | 135451 |
| MB17056 | 191569 | 157106 | 153905 | 155351 | 146281 | 136252 |
| MB17081 | 192056 | 158441 | 155914 | 156977 | 149902 | 137254 |
| MB17080 | 199543 | 157645 | 152764 | 155526 | 145146 | 137448 |
| MB17057 | 188453 | 154131 | 150438 | 152152 | 143230 | 137452 |
| MB17093 | 195807 | 160979 | 157252 | 159058 | 149234 | 137646 |
| MB17073 | 187906 | 155094 | 151940 | 153411 | 145848 | 137893 |
| MB17051 | 198862 | 164021 | 160914 | 162240 | 154624 | 138056 |
| MB17092 | 193078 | 158934 | 155322 | 157022 | 147570 | 138112 |
| MB17078 | 194868 | 157732 | 152508 | 155463 | 144991 | 139225 |
| MB17043 | 193051 | 160163 | 157638 | 158615 | 150842 | 139282 |
| MB17082 | 202218 | 163290 | 158657 | 161050 | 150067 | 142975 |
| MB17096 | 207652 | 168566 | 165394 | 166562 | 156679 | 143911 |
| MB17045 | 191882 | 160135 | 157190 | 158602 | 151395 | 144044 |
| MB17097 | 198707 | 162450 | 158934 | 160512 | 151448 | 144901 |
| MB17079 | 199205 | 160454 | 156515 | 158632 | 150954 | 145648 |
| MB17075 | 206904 | 167892 | 163984 | 166015 | 156259 | 145692 |
| MB17061 | 205434 | 167177 | 163288 | 165326 | 155941 | 146142 |
| MB17054 | 216080 | 175190 | 171968 | 173432 | 163969 | 147488 |
| MB17041 | 205535 | 165792 | 161108 | 163759 | 153714 | 147506 |
| MB17053 | 204928 | 168421 | 164296 | 166425 | 155034 | 147578 |
| MB17064 | 208836 | 170448 | 166606 | 168481 | 158842 | 147668 |
| MB17067 | 215352 | 173320 | 170278 | 171461 | 161503 | 147678 |
| MB17059 | 213035 | 172316 | 169264 | 170503 | 161090 | 147907 |
| MB17044 | 206910 | 168662 | 164467 | 166494 | 156705 | 148064 |
| MB17058 | 221285 | 184057 | 181235 | 182544 | 172926 | 152843 |
| MB17042 | 214336 | 173569 | 168171 | 171035 | 159865 | 154630 |
| MB17047 | 218450 | 177011 | 172184 | 174512 | 163658 | 156921 |
| MB17052 | 224349 | 186093 | 182479 | 184206 | 174900 | 160908 |
